## Supplementary Material for "A randomization-based causal inference framework for uncovering environmental exposure effects on human gut microbiota"

[1]Department of Statistics, Harvard University, Cambridge, MA, USA
[2]Institute for Medical Information Processing, Biometry, and Epidemiology, Faculty of Medicine, Ludwig-Maximilians-University München, Munich, Germany
[3]Institute of Epidemiology, Helmholtz Zentrum München, Neuherberg, Germany
[4]Department of Environmental Health, Harvard T. H. Chan School of Public Health, Boston, MA, USA
[5]Research Unit of Molecular Epidemiology, Helmholtz Zentrum München, Neuherberg, Germany
[6]German Center for Diabetes Research (DZD), München-Neuherberg, Germany
[7]ZIEL - Institute for Food & Health, Technical University of Munich, Freising, Germany
[8]Chair of Nutrition and Immunology, Technical University of Munich, Freising, Germany
[9]Institute of Computational Biology, Helmholtz Zentrum München, Neuherberg, Germany
[10]Department of Statistics, Ludwig-Maximilians-University München, Munich, Germany
[11]Center for Computational Mathematics, Flatiron Institute, New York, NY, USA
[12]Biostatistics Center, Massachusetts General Hospital and Harvard Medical School, Boston, MA, USA

-

#### Gut microbiome data description (Amplicon Sequence Variants)

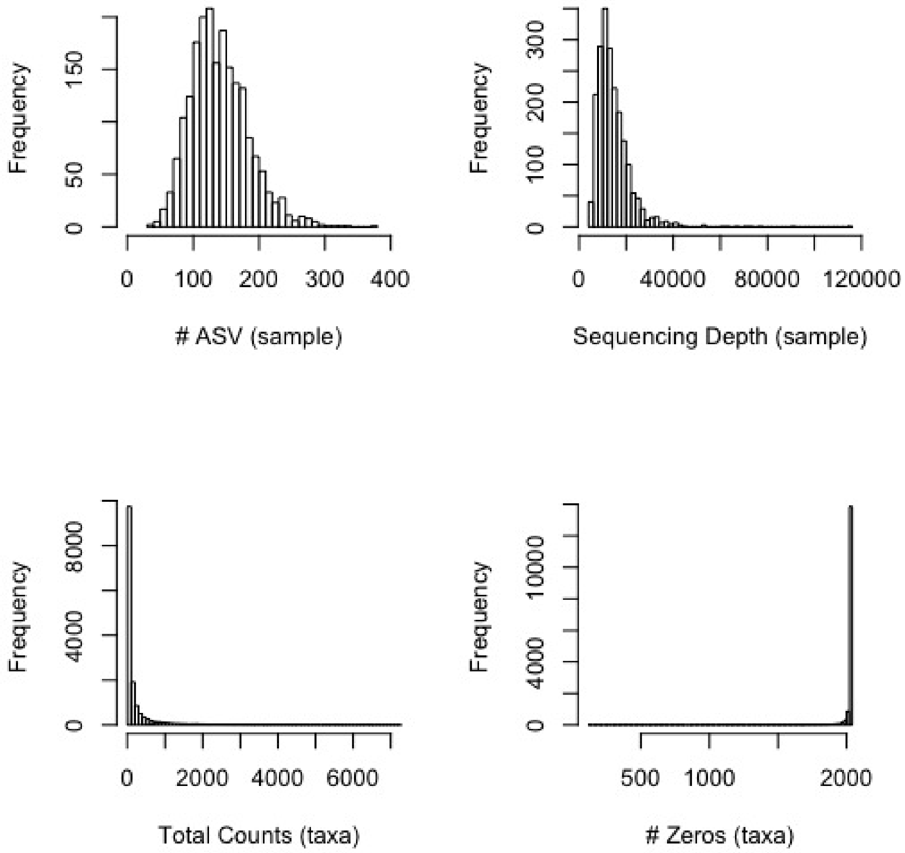


Figure 1: Gut microbiome data description. Number of observed ASV per sample (top left), sequencing depth per sample (top right), number of sequences per ASV (bottom left), number of zero count per ASV (bottom right).

| | Min. | $1^{st}$ Qu. | Median | Mean | $3^{rd}$ Qu. | Max. |
|---|---|---|---|---|---|---|
| nb. ASV (per sample) | 31 | 109 | 135 | 140 | 168 | 371 |
| nb. counts (per sample) | 4,696 | 9,696 | 12,716 | 14,470 | 17,292 | 115,055 |
| nb. counts (per taxa) | 1 | 16 | 61 | 1,863 | 219 | 729,636 |
| nb. zeros (per taxa) | 122 | 2,030 | 2,033 | 2,016 | 2,033 | 2,033 |

Table 1: Gut microbiome data description. Number of observed ASV per sample, sequencing depth per sample, number of sequences per ASV, number of zero count per ASV.

### Balance diagnostics for matching

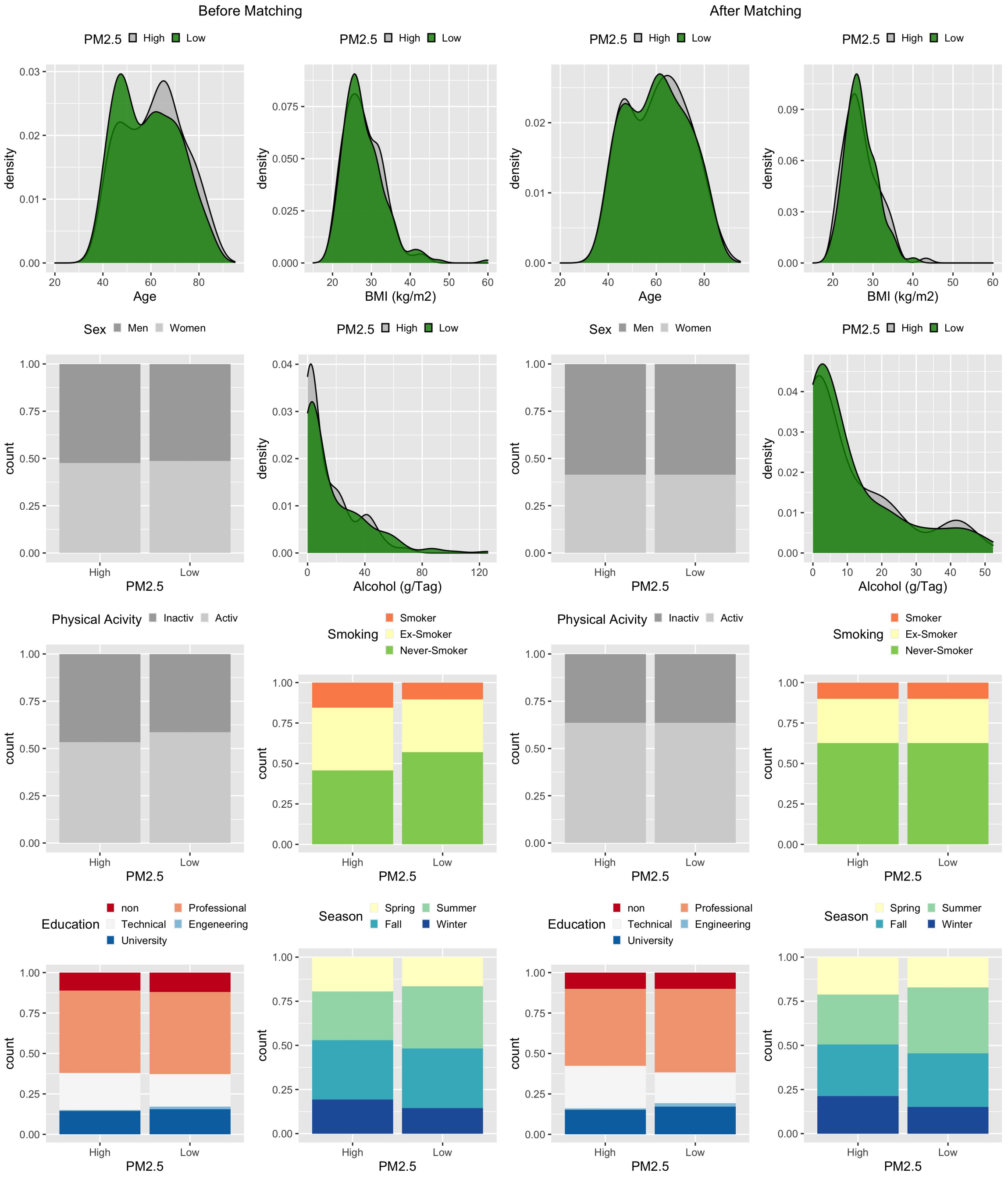


Figure 2: Empirical distributions of the matched covariates among the subjects under the intervention vs. not in the original (left panel) and the balanced (right panel) data for the air pollution reduction hypothetical experiment.

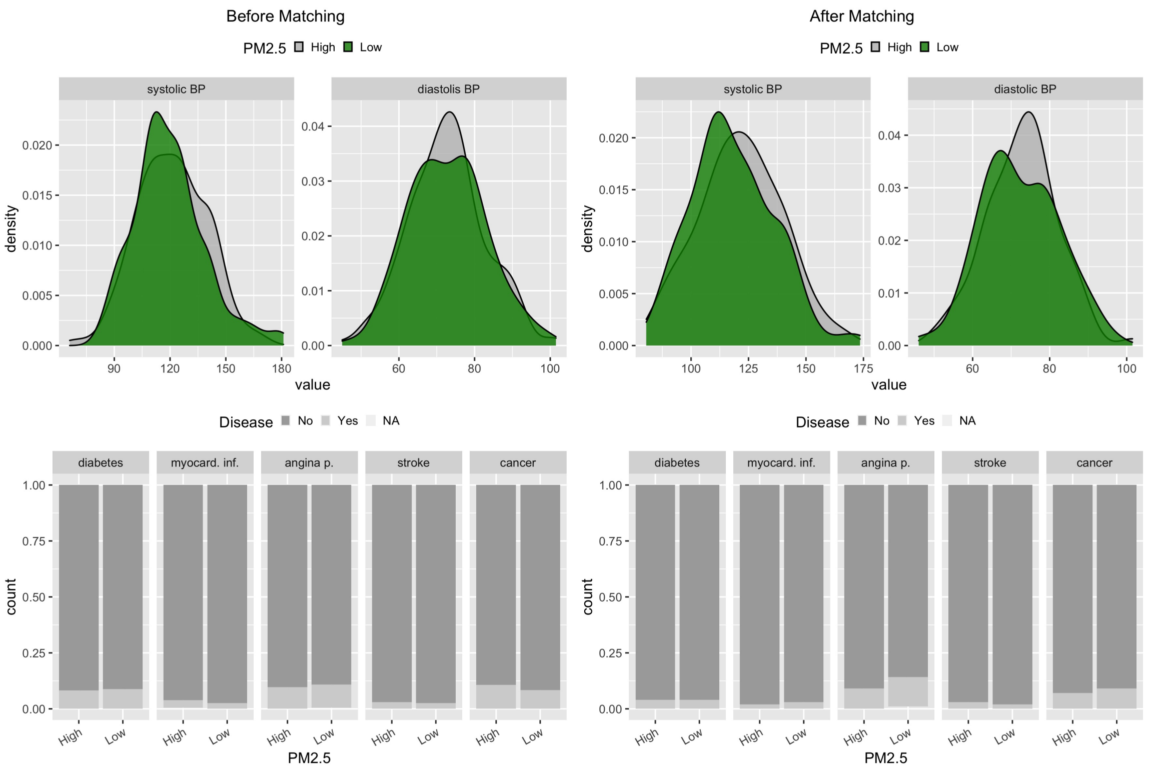


Figure 3: Empirical distributions of the disease covariates among the subjects under the intervention vs. not in the original (left panel) and the balanced (right panel) data for the air pollution reduction hypothetical experiment.

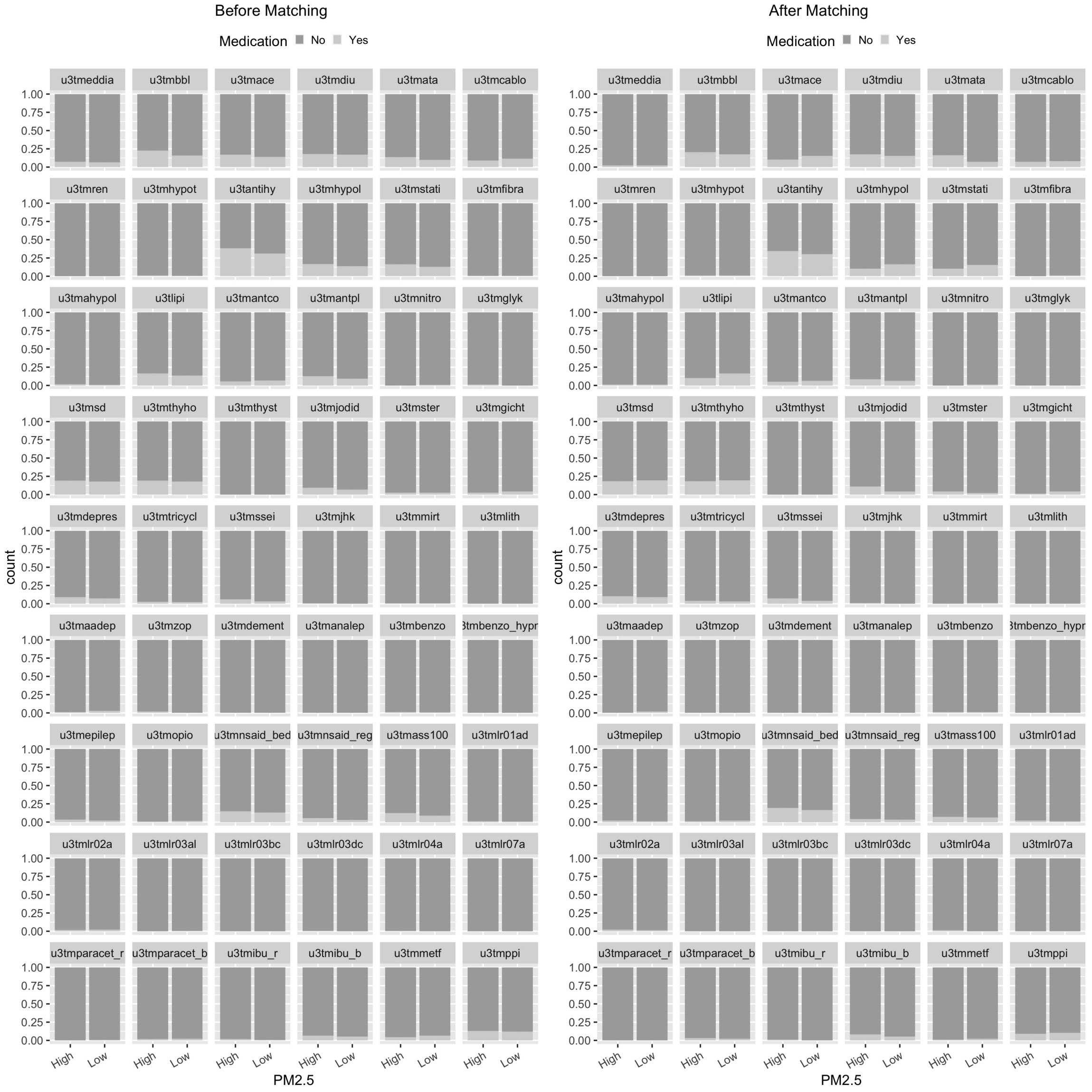


Figure 4: Empirical distributions of the medication covariates among the subjects under the intervention vs. not in the original (left panel) and the balanced (right panel) data for the air pollution reduction hypothetical experiment.

### Smoking

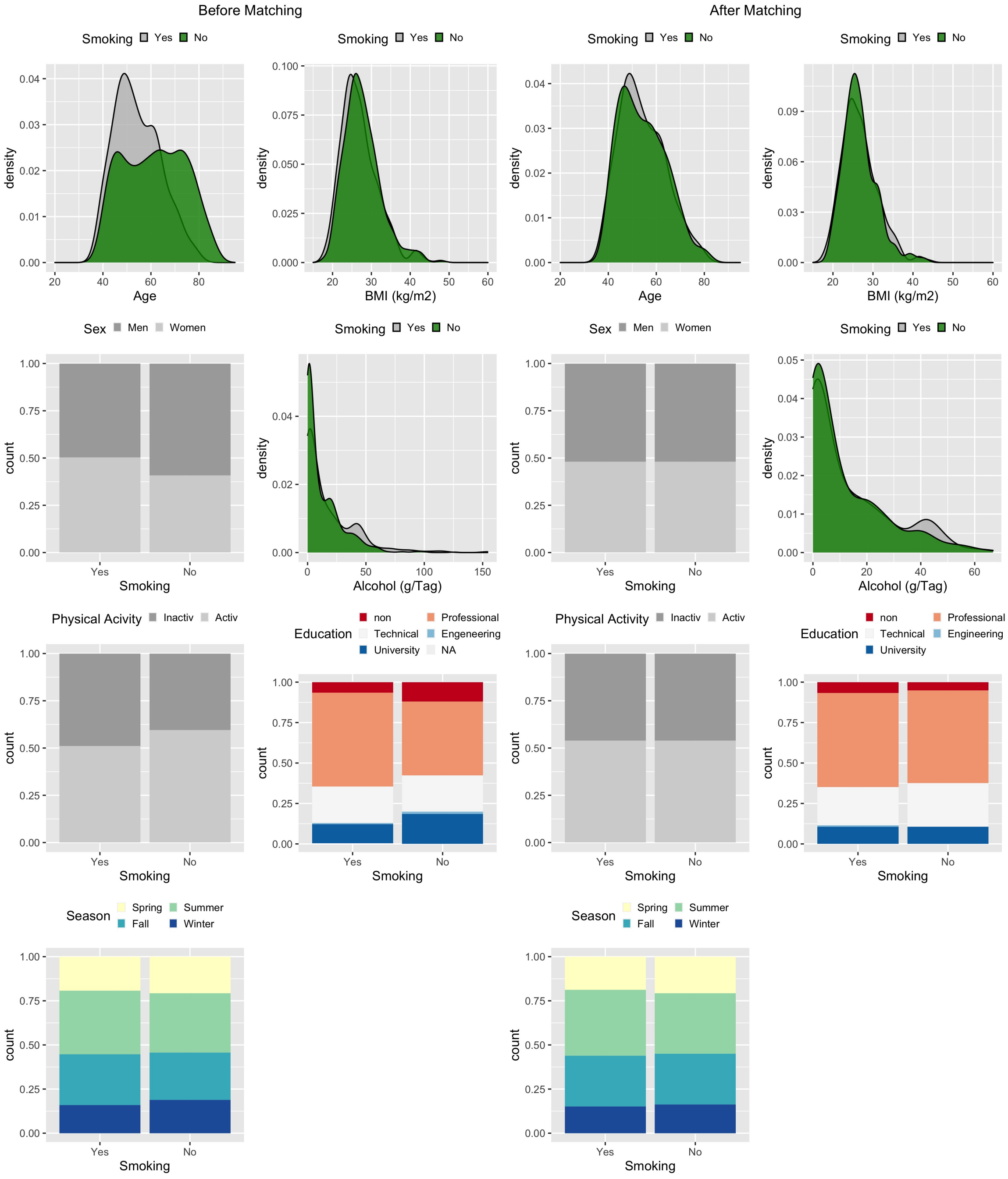


Figure 5: Empirical distributions of the matched covariates among the subjects under the intervention vs. not in the original (left panel) and the balanced (right panel) data for the smoking prevention hypothetical experiment.

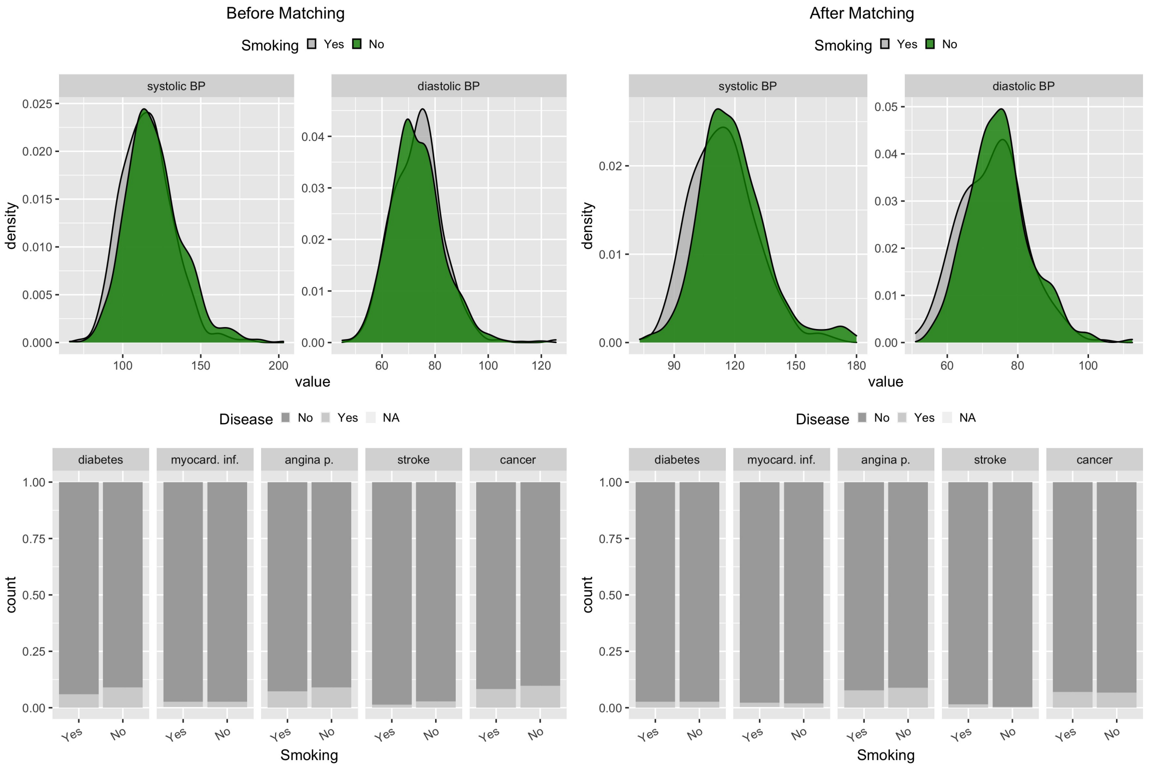


Figure 6: Empirical distributions of the diseases covariates among the subjects under the intervention vs. not in the original (left panel) and the balanced (right panel) data for the smoking prevention hypothetical experiment.

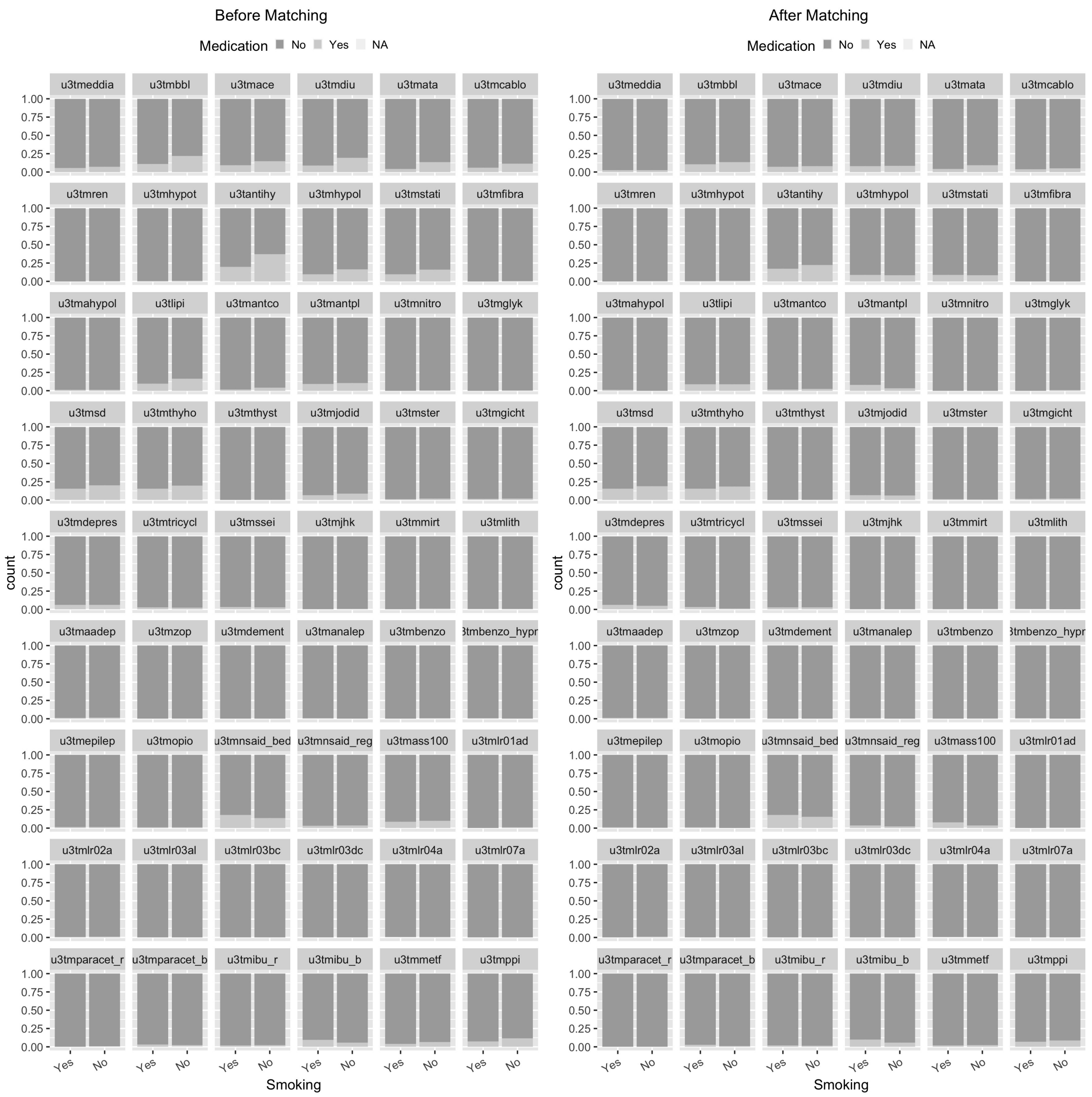


Figure 7: Empirical distributions of the medication covariates among the subjects under the intervention vs. not in the original (left panel) and the balanced (right panel) data for the smoking prevention hypothetical experiment.

### Balance diagnostics for nutrition covariates after matching

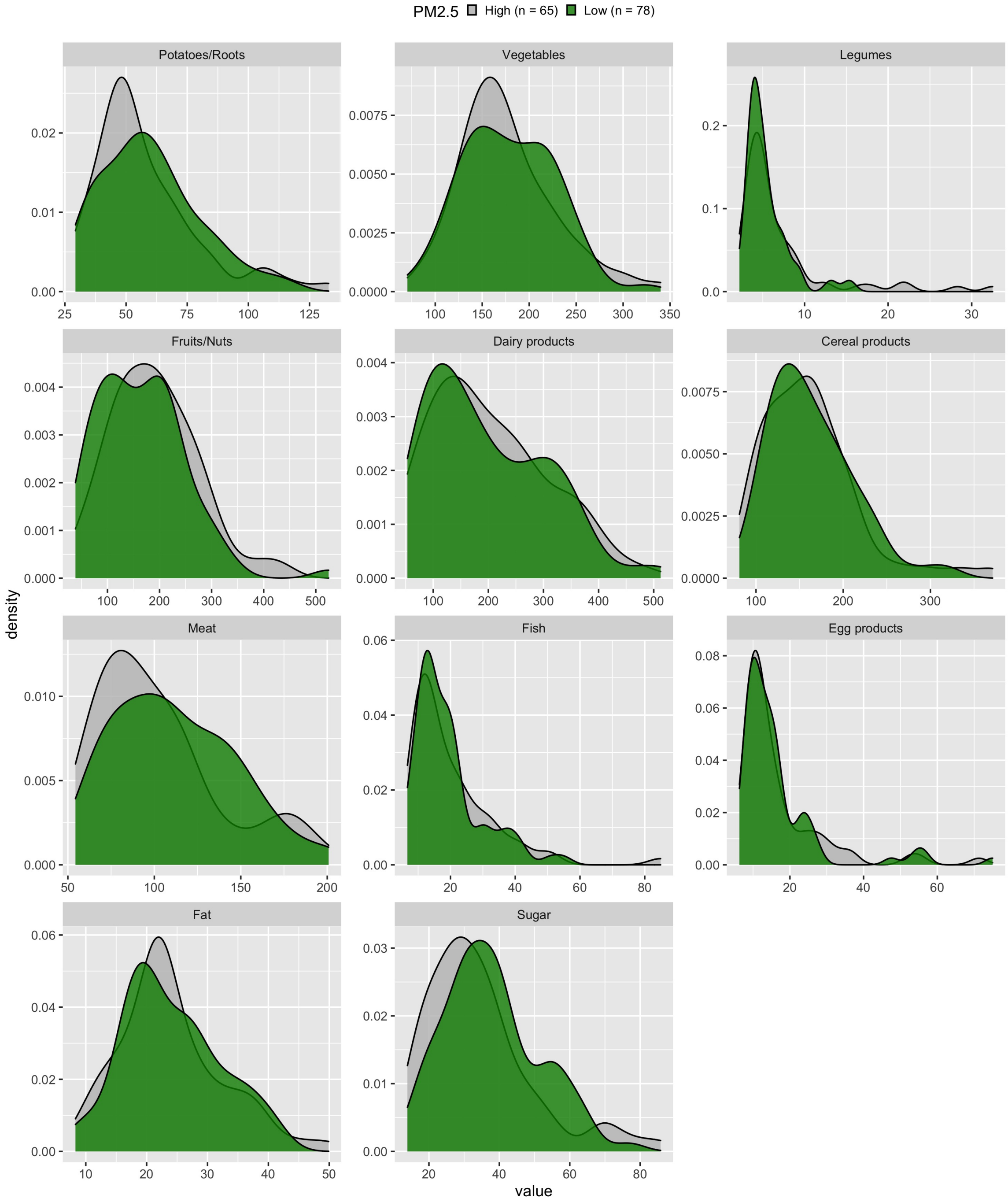


Figure 8: Empirical distributions of the nutrition covariates among the subjects under the intervention vs. not in the balanced data for the air pollution reduction hypothetical experiment.

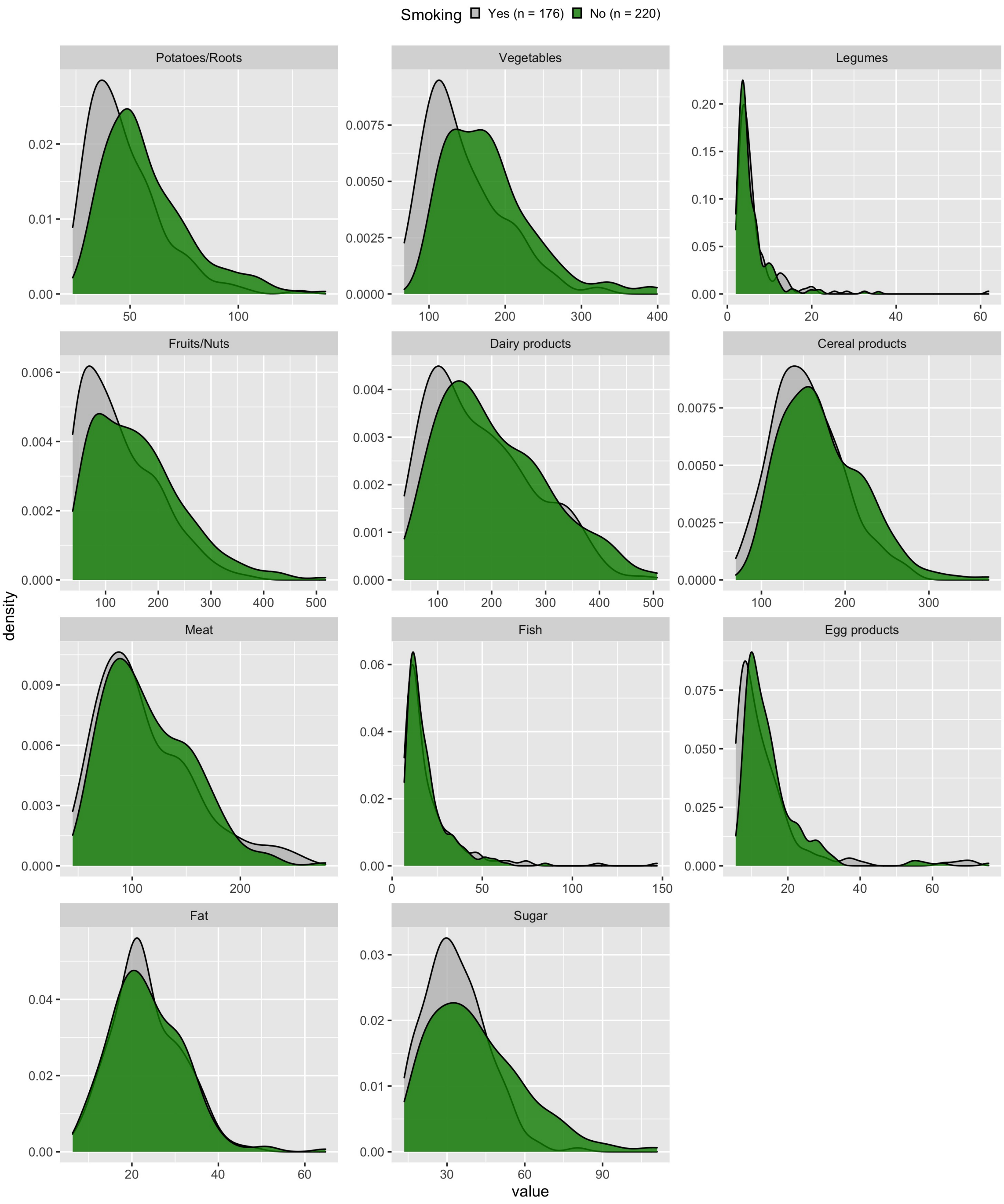


Figure 9: Empirical distributions of the nutrition covariates among the subjects under the intervention vs. not in the balanced data for the smoking prevention hypothetical experiment.

#### Comparison of permutation and asymptotic null randomization distribution

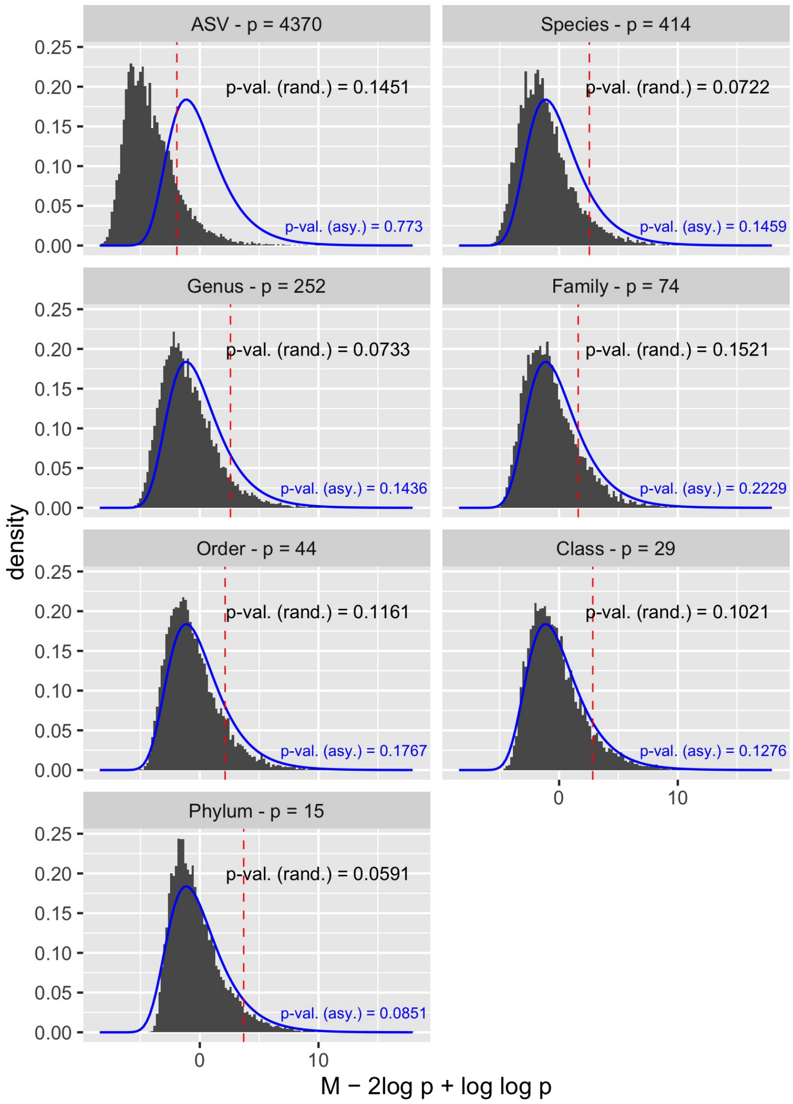


Figure 10: Permutation-based (grey) and asymptotic (blue) null randomization distributions for the air pollution reduction hypothetical experiment.

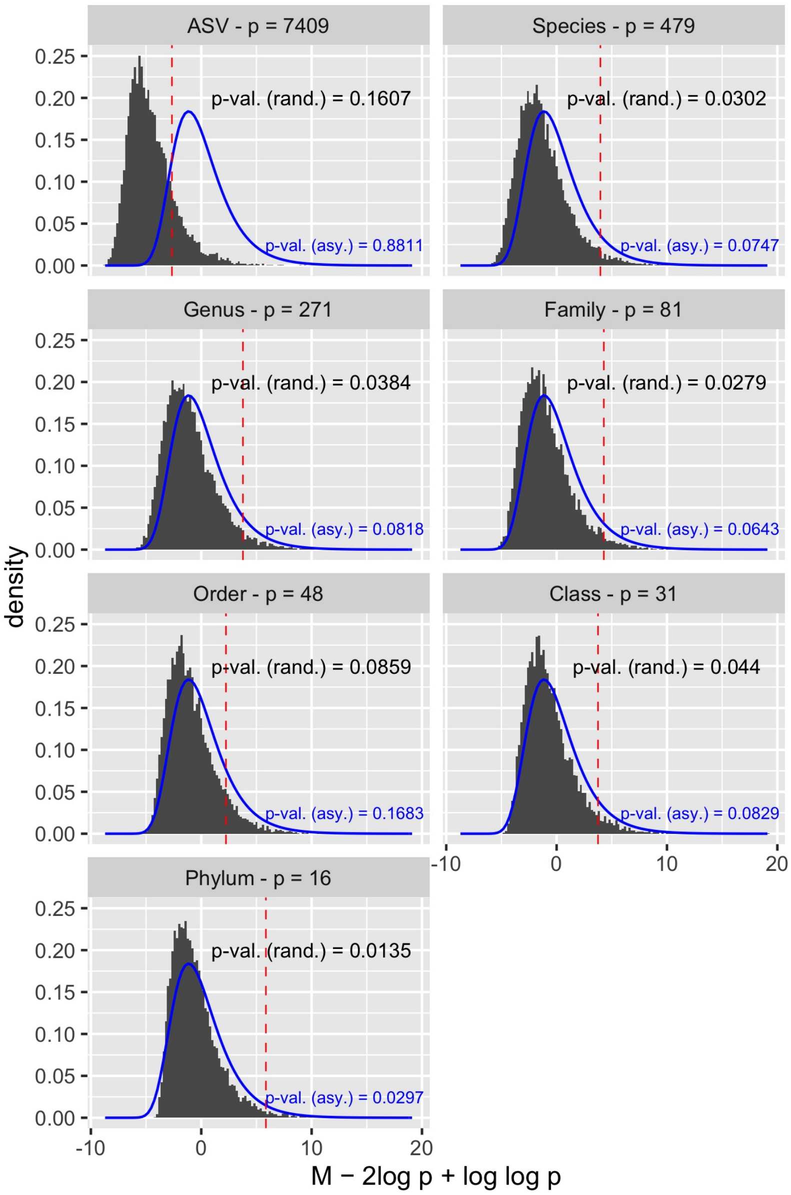


Figure 11: Permutation-based (grey) and asymptotic (blue) null randomization distributions for the smoking prevention hypothetical experiment.

#### Reference selection for DACOMP

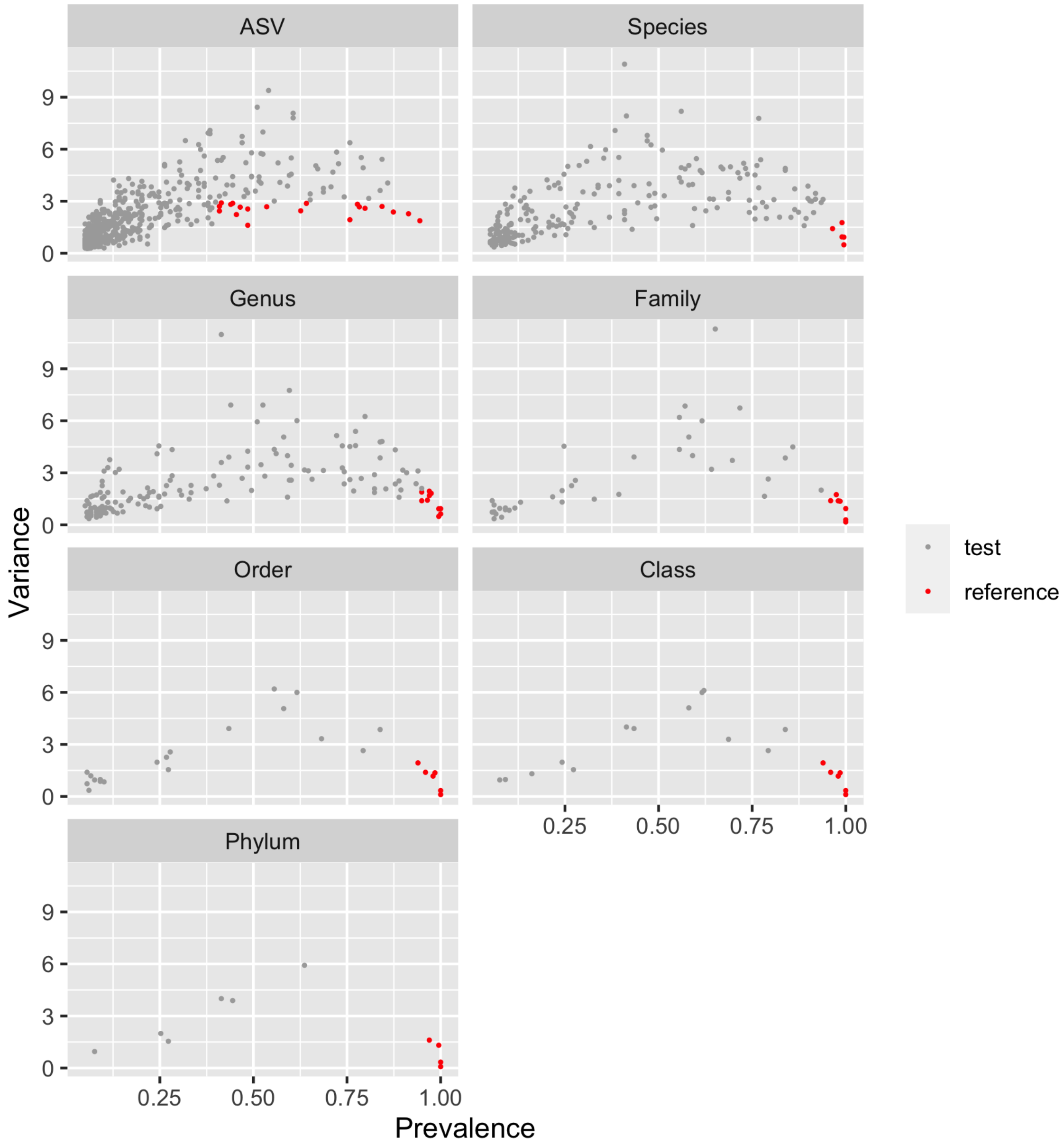


Figure 12: Reference set selection in the air pollution reduction experiment. A taxa enters the set $R = (r_1, ..., r_F)$ if it has low variance ($< 2$) and high prevalence ($> 90\%$). For the analyses at the ASV level, we chose the variance to be $< 3$ and the prevalence to be $> 40\%$ as thresholds in order the have at least one reference per subject.

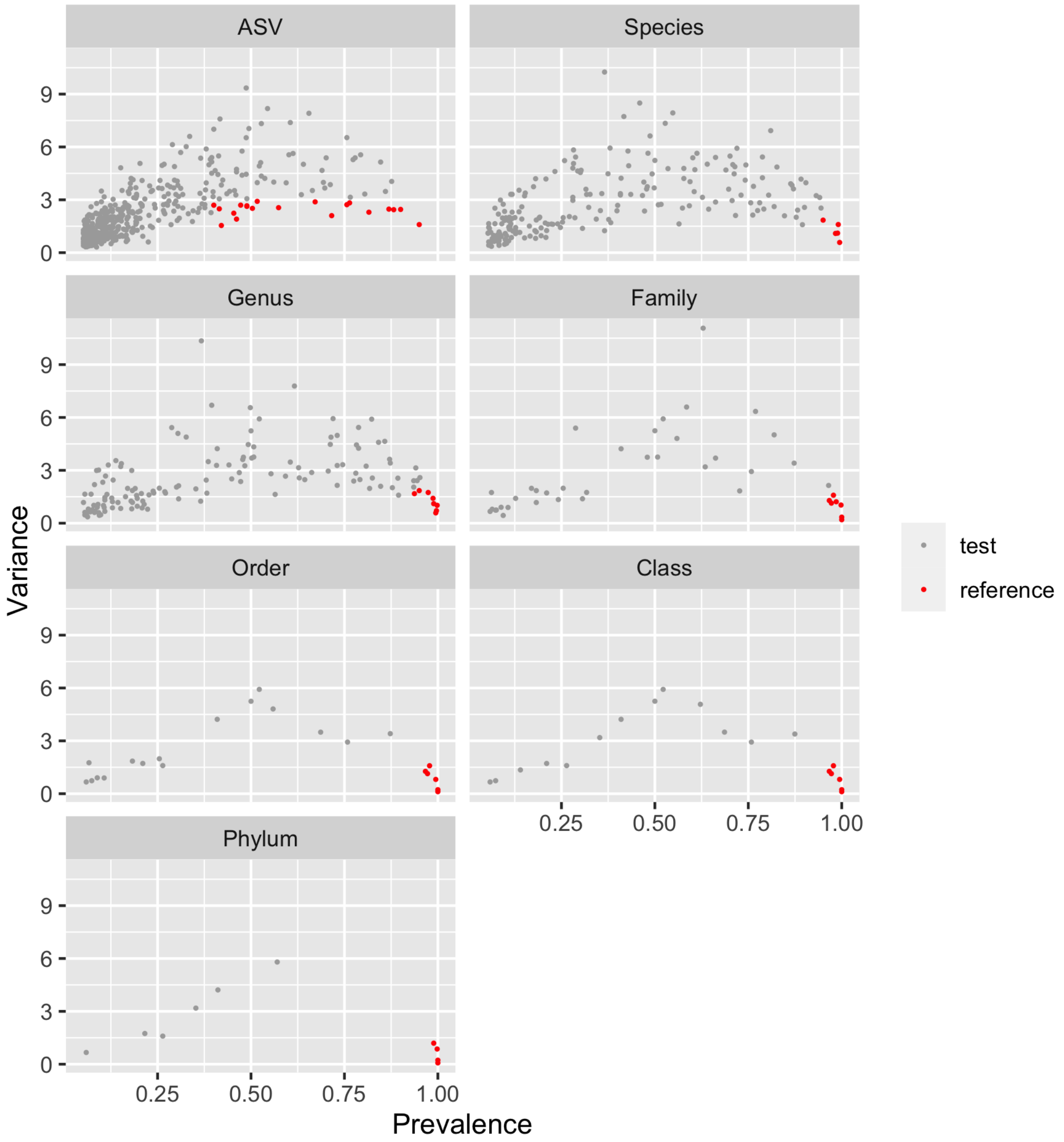


Figure 13: Reference set selection in the smoking prevention experiment. A taxa enters the set $R = (r_1, ..., r_F)$ if it has low variance ($< 2$) and high prevalence ($> 90\%$). For the analyses at the ASV level, we chose the variance to be $< 3$ and the prevalence to be $> 40\%$ as thresholds in order the have at least one reference per subject.

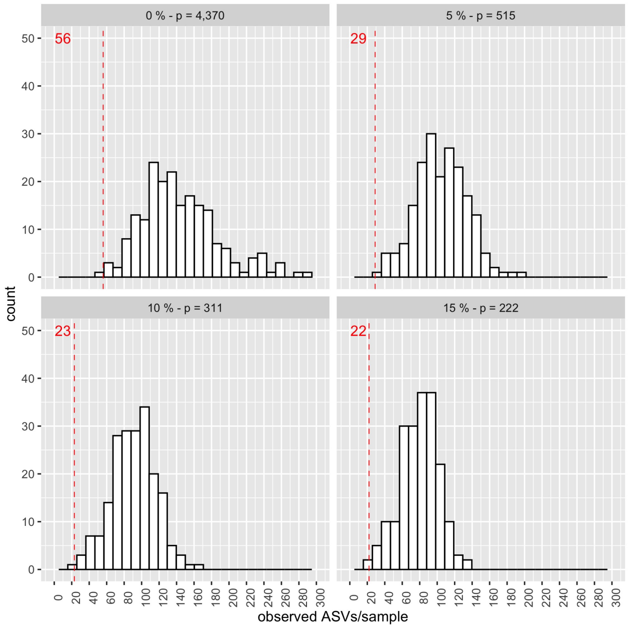


Figure 14: Distribution of number of ASVs per sample when data is filtered at different ASV prevalence thresholds (0%, 5%, 10%, 15%) in the air pollution reduction experiment. Red value: minimum observed ASVs per sample.

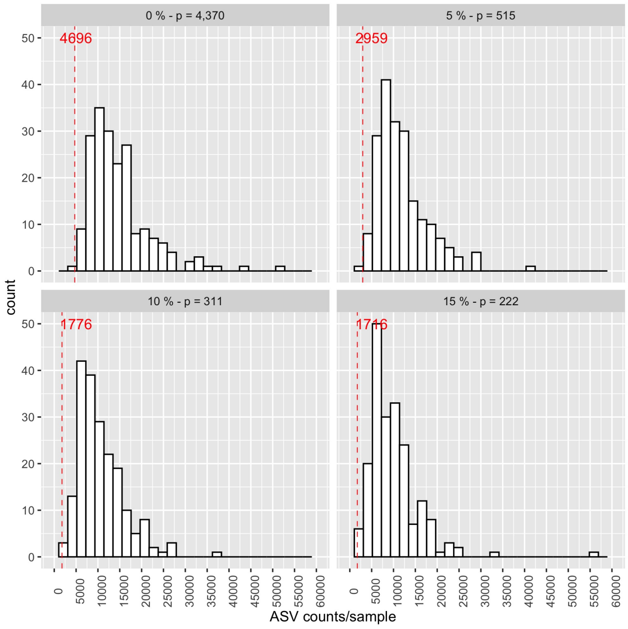


Figure 15: Distribution of the total ASV counts per sample when data is filtered at different ASV prevalence thresholds (0%, 5%, 10%, 15%) in the air pollution reduction experiment. Red value: minimum ASV counts per sample.

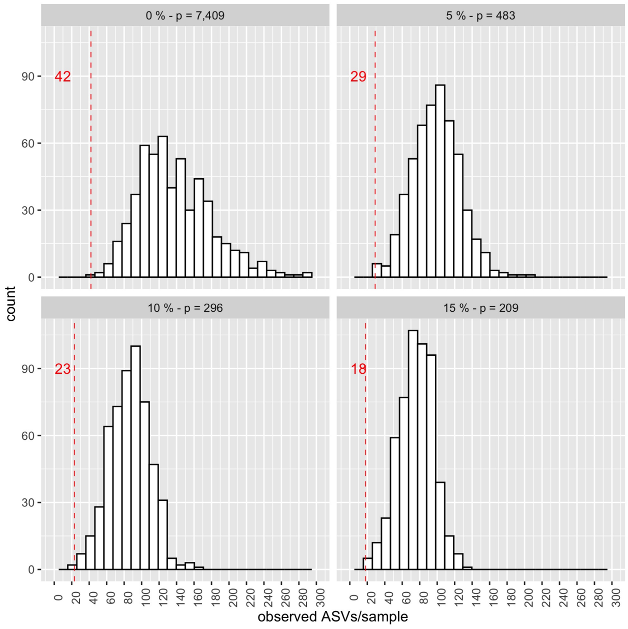


Figure 16: Distribution of number of ASVs per sample when data is filtered at different ASV prevalence thresholds (0%, 5%, 10%, 15%) in the smoking prevention reduction experiment. Red value: minimum observed ASVs per sample.

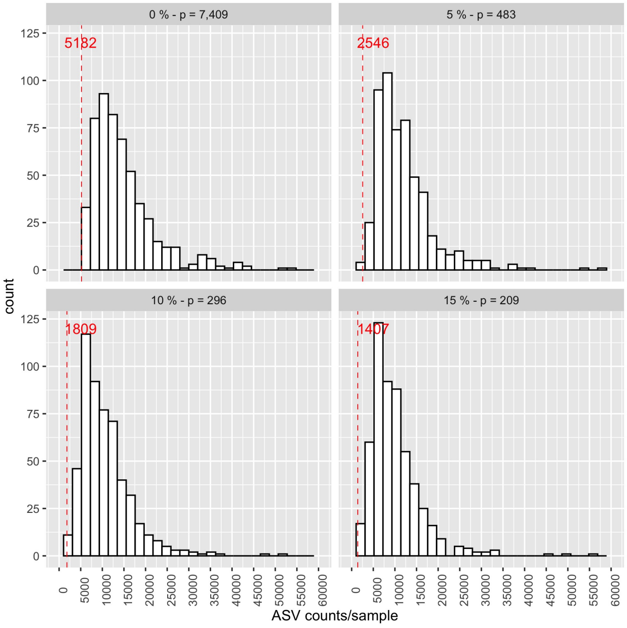


Figure 17: Distribution of the total ASV counts per sample when data is filtered at different ASV prevalence thresholds (0%, 5%, 10%, 15%) in the smoking prevention experiment. Red value: minimum ASV counts per sample.

| | Kingdom | Phylum | Class | Order | Family | Genus | Species | p-value$_{adj}$ |
|---|---|---|---|---|---|---|---|---|
| ASV p = 515 | Bacteria | Firmicutes | Clostridia | Clostridiales | Ruminococcaceae | Anaerotruncus | NA | 0.1461 (+) |
| | Bacteria | Firmicutes | Clostridia | Clostridiales | Ruminococcaceae | Anaerotruncus | NA | 0.0362 (-) |
| **Species** p = 220 | Bacteria | Firmicutes | Clostridia | Clostridiales | Lachnospiraceae | Blautia | faecis | 0.0357 (+) |
| | Bacteria | Firmicutes | Clostridia | Clostridiales | Lachnospiraceae | Marvinbryantia | NA | 0.0181 (+) |
| **Genus** p = 149 | Bacteria | Firmicutes | Clostridia | Clostridiales | Lachnospiraceae | Marvinbryantia | NA | 0.0120 (+) |
| **Family** p = 44 | | | | | | | | |
| **Order** p = 25 | | | | | | | | |
| **Class** p = 19 | | | | | | | | |
| **Phylum** p = 10 | | | | | | | | |

Table 2: Air pollutiion reduction experiment results. Differentially abundant bacteria and adjusted Fisher p-values for 10,000 iterations at 5% prevalence filtering. Selected adjusted p-values $\leq 0.2$ (sign of abundance difference: y(1) - y(0)).

| | Kingdom | Phylum | Class | Order | Family | Genus | Species | p-value$_{adj}$ |
|---|---|---|---|---|---|---|---|---|
| **ASV** p = 483 | Bacteria | Firmicutes | Clostridia | Clostridiales | Ruminococcaceae | Ruminococcus-1 | NA | 0.1250 (+) |
| | Bacteria | Firmicutes | Clostridia | Clostridiales | Ruminococcaceae | Ruminococcaceae-UCG-002 | NA | 0.1458 (+) |
| | Bacteria | Firmicutes | Clostridia | Clostridiales | Lachnospiraceae | Lachnospiraceae-NK4A136-group | NA | 0.1124 (+) |
| **Species** | Bacteria | Firmicutes | Clostridia | Clostridiales | Christensenellaceae | Christensenellaceae-R-7-group | NA | 0.0201 (+) |
| p = 211 | Bacteria | Firmicutes | Clostridia | Clostridiales | Ruminococcaceae | Ruminococcaceae-UCG-005 | NA | 0.1124 (+) |
| | Bacteria | Firmicutes | Clostridia | Clostridiales | Ruminococcaceae | Ruminococcaceae-UCG-003 | NA | 0.1297 (+) |
| | Bacteria | Firmicutes | Clostridia | Clostridiales | Lachnospiraceae | Coprococcus-1 | catus | 0.0392 (+) |
| | Bacteria | Tenericutes | Mollicutes | NB1-n | NA | NA | NA | 0.1458 (+) |
| | Bacteria | Tenericutes | Mollicutes | Mollicutes-RF9 | NA | NA | NA | 0.1791 (+) |
| | Bacteria | Firmicutes | Clostridia | Clostridiales | Ruminococcaceae | Ruminococcaceae-UCG-002 | NA | 0.1476 (+) |
| | Bacteria | Firmicutes | Clostridia | Clostridiales | Ruminococcaceae | Ruminococcaceae-UCG-003 | NA | 0.0127 (+) |
| | Bacteria | Firmicutes | Clostridia | Clostridiales | Ruminococcaceae | Ruminococcaceae-UCG-005 | NA | 0.1975 (+) |
| | Bacteria | Firmicutes | Clostridia | Clostridiales | Ruminococcaceae | Ruminococcus-1 | NA | 0.1691 (+) |
| **Genus** | Bacteria | Firmicutes | Clostridia | Clostridiales | Ruminococcaceae | Ruminococcaceae-NK4A214-group | NA | 0.1476 (+) |
| p = 140 | Bacteria | Firmicutes | Clostridia | Clostridiales | Christensenellaceae | Christensenellaceae-R-7-group | NA | 0.0611 (+) |
| | Bacteria | Firmicutes | Clostridia | Clostridiales | Lachnospiraceae | Lachnospira | NA | 0.0377 (+) |
| | Bacteria | Firmicutes | Clostridia | Clostridiales | Lachnospiraceae | Lachnospiraceae-NK4A136-group | NA | 0.1781 (+) |
| | Bacteria | Firmicutes | Clostridia | Clostridiales | Lachnospiraceae | Coprococcus-1 | NA | 0.0611 (+) |
| | Bacteria | Tenericutes | Mollicutes | NB1-n | NA | NA | NA | 0.1882 (+) |
| | Bacteria | Tenericutes | Mollicutes | Mollicutes-RF9 | NA | NA | NA | 0.1166 (+) |
| | Bacteria | Firmicutes | Clostridia | Clostridiales | Christensenellaceae | NA | NA | 0.0199 (+) |
| **Family** | Bacteria | Tenericutes | Mollicutes | NB1-n | NA | NA | NA | 0.0450 (+) |
| p = 41 | Bacteria | Tenericutes | Mollicutes | Mollicutes-RF9 | NA | NA | NA | 0.0512 (+) |
| **Order** | Bacteria | Tenericutes | Mollicutes | NB1-n | NA | NA | NA | 0.0375 (+) |
| p = 22 | Bacteria | Tenericutes | Mollicutes | Mollicutes-RF9 | NA | NA | NA | 0.0404 (+) |
| **Class** p = 19 | Bacteria | Tenericutes | Mollicutes | NA | NA | NA | NA | 0.0039 (+) |
| **Phylum** p = 10 | Bacteria | Tenericutes | NA | NA | NA | NA | NA | 0.0018 (+) |

Table 3: Smoking prevention experiment results. Differentially abundant bacteria and adjusted Fisher p-values for 10,000 iterations at 5% prevalence filtering. Selected adjusted p-values $\leq 0.2$ (sign of abundance difference: y(1) - y(0)).

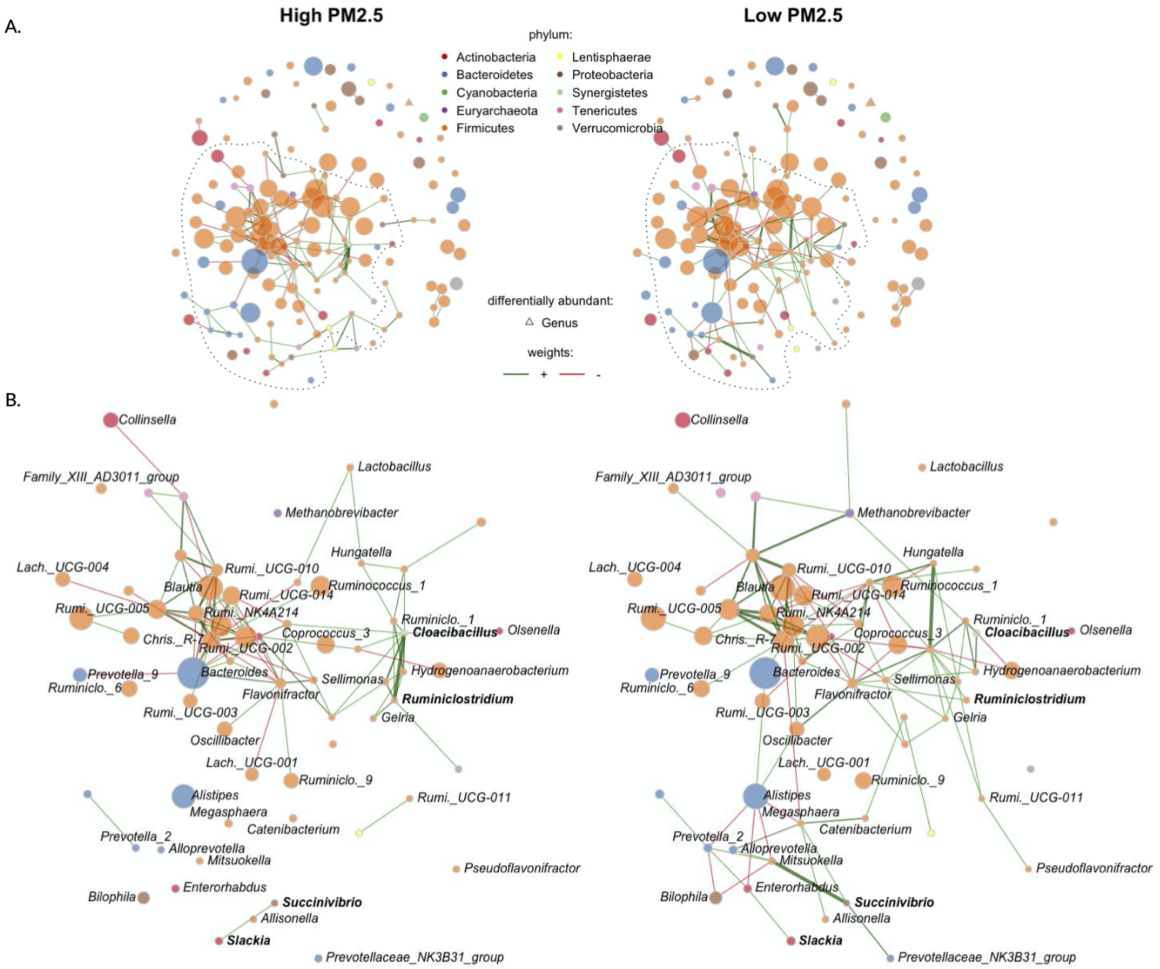


Figure 18: Genus-genus associations for subject under the air pollution reduction experiment vs. not (n = 99, p = 149). (A) Visualization of the between genera partial correlations estimated with the SPIEC-EASI method. Edges thickness is proportional to partial correlation, and color to direction: red: negative partial correlation, green: positive partial correlation. Node size is proportional to the centered log ratio of the genus abundances, and color is according to phyla. Triangle shaped nodes are differentially abundant (see Figure 3). (B) Zoom in largest connected component and differential associations (bold genera).

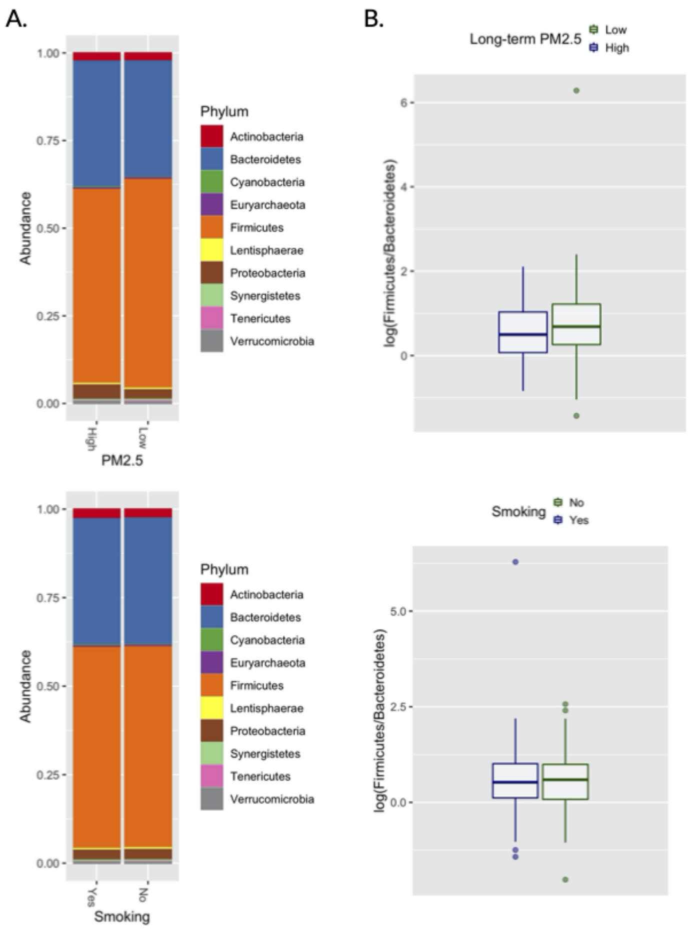


Figure 19: Phyla comparison.
